# Noise facilitates entrainment of a population of uncoupled limit cycle oscillators

**DOI:** 10.1101/2022.03.28.486061

**Authors:** Vojtech Kumpost, Lennart Hilbert, Ralf Mikut

**Affiliations:** Institute for Automation and Applied Informatics, Karlsruhe Institute of Technology, Eggenstein-Leopoldshafen, Germany; Institute of Biological and Chemical Systems - Biological Information Processing, Karlsruhe Institute of Technology, Eggenstein-Leopoldshafen, Germany; Zoological Institute, Department of Systems Biology and Bioinformatics, Karlsruhe Institute of Technology, Karlsruhe, Germany

## Abstract

Many biological oscillators share two properties: they are subject to stochastic fluctuations (noise) and they must reliably adjust their period to changing environmental conditions (entrainment). While noise seems to distort the ability of single oscillators to entrain, in populations of oscillators noise allows entrainment for a wider range of input amplitudes and periods. Here, we investigate, how this effect depends on the noise intensity and the number of oscillators in the population. We have found that, if a population consists of a sufficient number of oscillators, increasing noise intensity leads to faster entrainment after a phase change of the input signal (jet lag) and increases sensitivity to low-amplitude input signals.

**SIGNIFICANCE:** Live is characterized by rhythms, such as daily changes in activity or the heartbeat. These rhythms are reflected in molecular oscillations generated at the level of individual cells. These oscillations are inherently noisy, but still cells reliably synchronize to external signals and provide reliable timing for other biological processes. Here, we show how noise can be beneficial to cell populations in terms of synchronization to external signals. Specifically, noise can increase the sensitivity to weak external signals and speed up adjustment to jet-lag-like perturbations.

## INTRODUCTION

Many cellular processes show oscillatory behavior. This includes circadian clocks (1), cardiac pacemakers (2), and various signaling proteins like p53 (3) and NF-κB (4). Although the rhythms generated by these systems manifest on different timescales and have different functions within the organism, in terms of a mathematical representation, they can be modeled as a limit cycle oscillator that consists of a negative feedback loop with a delay (5). Furthermore, cellular oscillators constantly adjust their phase and period based on the changes in the environment, thereby achieving entrainment (6). In this way, cellular processes can be reliably timed with respect to environmental conditions. Cellular processes, including oscillators, are frequently affected by noise stemming from a low number of reacting molecules (7). In consequence, a low number of molecular interactions occur at stochastically distributed times, resulting in stochastic variations in the periodic gene expression. Although noise and entrainment are individually recognized to influence cellular oscillators, it remains unclear whether noise is generally detrimental to entrainment, or might in fact be co-opted in the entrainment of cellular oscillators.

The circadian clock is one of the most prominent biological oscillators and ensures the correct timing of various biological processes with respect to the time of the day (8). The circadian clock in mammals is organized hierarchically with a central peacemaker of highly coupled oscillators in the suprachiasmatic nucleus (SCN) and weakly coupled or uncoupled oscillators in the peripheral tissues (9). While a lot of research focused on the SCN, the limited coupling in peripheral tissues results in distinctly different dynamics in the population of peripheral tissue oscillators. For example, the weaker coupling between peripheral clocks allows entrainment for a wider range of input periods (10) and additional external cues such as changes in ambient temperature (11). The population of uncoupled oscillators is also relevant in the correct analysis of the circadian bioluminescence assays (12, 13). Here, a better understanding of the dynamics at the population level in the presence of noise would help in inferring the single-cell behavior from the population-level recordings. For those reasons, it is of great interest to better understand the significance of noise in the entrainment of uncoupled stochastic oscillators.

The implications of noise on the population-level entrainment are, however, still poorly understood. In this study, we examine the effect of noise on a population of uncoupled stochastic oscillators for different population sizes (number of oscillators), noise intensities, and amplitudes and periods of the input signal. Previous work found that noise allows population-level entrainment to a wider range of input amplitudes and periods than a single stochastic or deterministic oscillator (14). We extend those findings by examining the change in the entrainment for varying population size and noise intensity. In addition, we examine how different levels of noise influence the response to a perturbing pulse under constant pacing conditions (phase response curve) and the recovery from a phase shift in the input signal (jet lag). We have found that noise expands the range of input amplitudes and periods for which entrainment occurs, with an optimal noise intensity for a given number of oscillators in the population. In our simulations, noise also increases the response of the oscillator population to a perturbation and shortens jet lag. Finally, we used the canonical Van der Pol model, which represents the class of limit cycle oscillators, to show that we come to the same conclusions. Thus these findings might be interesting not only for the modeling of cellular oscillators but also for the wider class of natural and bio-inspired technical systems with uncoupled or loosely coupled oscillators under the influence of external triggers and noise.

## METHODS

### Kim-Forger model

The scaled minimal Kim-Forger model (15) is

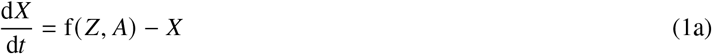

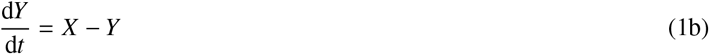

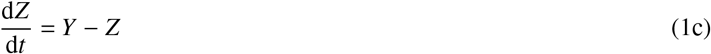

where f(*Z, A*) is a Kim-Forger function (15, 16) that has form

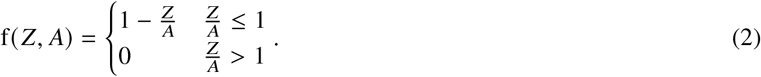

The only free parameter we set to *A* = 0.1, which corresponds to a limit cycle oscillator with a maximal possible amplitude (SI Figure 1). Further, for computational convenience, we scaled the model with a parameter *τ* = 3.66 so the free-running period is 1.

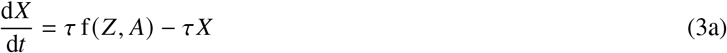

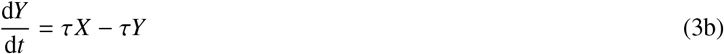

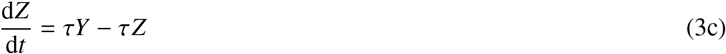

Next, we extended the model by an additive light input *I* as done before to model the entrainment in zebrafish cell cultures (17)

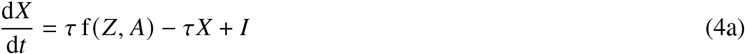

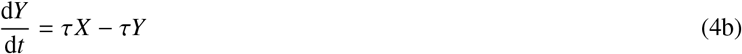

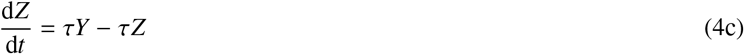

The molecular noise is estimated using stochastic differential equations (18)

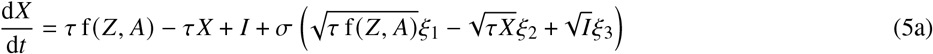

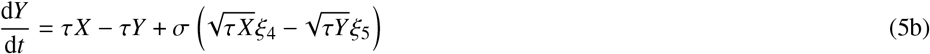

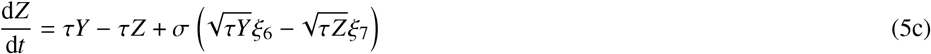

where *ξ*_*i*_ are independent Wiener processes and *σ* is noise intensity. The main disadvantage of this type of equations is that for high noise intensities the concentrations might reach negative values, which make the evaluation of the terms under the square roots impossible in the real domain (19). In the most of this study we used *σ* low enough to avoid this problem. For the few cases when we needed higher noise intensities, we implemented also the Gillespie method (20, 21) for the chemical reactions that are represented by Eq. (5).

The SDE model was simulated using the Euler-Maruyama method with an integration step dt = 0.0001. The integration step is sufficiently low to give comparable results with an accurate deterministic adaptive method for *σ* = 0 (SI Figure 2) as well as the exact Gillespie method for higher values of *σ* (SI Figure 3, SI Figure 4).

A population of uncoupled oscillators was simulated by calculating a mean of a repeated numerical simulation as

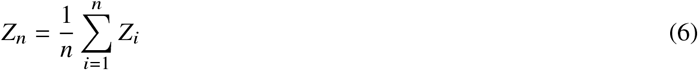

where *n* is the number of oscillators in the population.

### Van der Pol model

As a generic model, we used the Van der Pol model in form with an external input and noise (22)

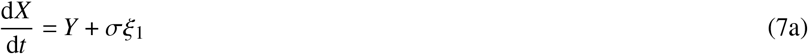

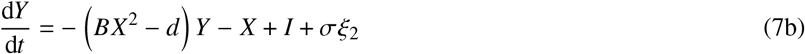

where *σ* is noise intensity, *ξ*_*i*_ are independent Wiener processes, *I* is input, and *d* and *B* are free parameters that we set according to the previous study (22) to represent a limit cycle oscillator (*d* = 2, *B* = 10) and a noise-driven oscillator (*d* = − 0.1 and *B* = 1).

Further, for computational convenience, we scaled the time with a parameter *τ* so the free-running period of the deterministic model is 1. The final equations read

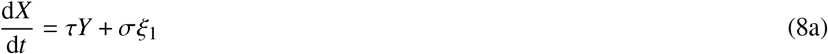

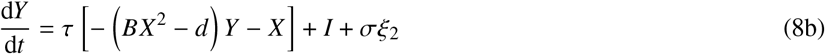

where *τ* = 7.63 for the limit cycle oscillator and *τ* = 6.2 for the noise-driven oscillator.

### Model input, entrainment, entrainment area

We consider a square input as

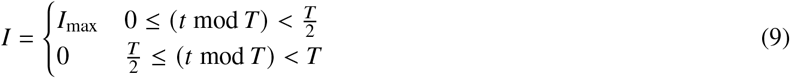

where *T* is the input period and *I*_max_ is the input amplitude. The output period of the model is estimated as the position of the highest peak of its autocorrelation function (*T*_out_) calculate for lags up to time 10, which corresponds to time series shown in SI Figures 5, 6, 7. The model is considered entrained if

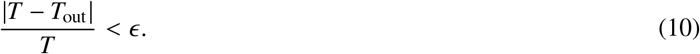

We set *∈* = 0.02. Examples of entrained and unentrained traces are in SI Figures 5, 6, and 7. To increase the efficiency of the Arnold tongue estimation we used a binary search for the border between the entrained and unentrained areas (14). With this numerical method the search is initialized with two points representing the minimal and maximal input amplitudes of interest. The minimal amplitude is set as unentrained and the maximal amplitude is set as entrained. At each step the middle point between the maximal and minimal amplitude is evaluated and assigned as entrained or unentrained according to Eq. (10).

The search is stopped when the difference between the entrained and unentrained amplitude is less than 0.01. Repeating this procedure we obtain the border amplitudes (*b*) for a series of input periods. The entrainment area is then defined as

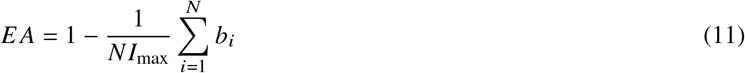

where *N* is the number of periods for which the entrainment border was estimated and *I*_max_ is the maximal amplitude that was considered in the search.

### Software implementation

All numerical experiments were implemented in Julia and are freely available as a GitHub repository at https://github.com/vkumpost/stoosc.

## RESULTS

### Model of a population of uncoupled stochastic oscillators

We wish to examine how molecular noise affects the entrainment of a population of cellular oscillators. For this purpose, we use a minimal Kim-Forger model (15) with additive light input and multiplicative noise terms, which represent the molecular noise stemming from the discrete nature of cellular chemical interactions (18). We have shown previously that such a simple structure captures the entrainment dynamics in a circadian bioluminescence reporter assay with high precision while reducing the number of adjustable model parameters (17). The core model consists of three variables connected in a negative feedback loop and represents the dynamics of one cellular oscillator (Figure 1A). The simplified Kim-Forger model (see Methods) has one free parameter for which we assigned a fixed value so as to obtain a limit cycle oscillator with high-amplitude oscillations (SI Figure 1). For convenience, the model is scaled to oscillate with a unit period if noise terms and input signals are set to zero (see Methods). To obtain a population output from the single-cell model, we repeat the simulation *n* times and calculate the mean over the individual trajectories, thereby representing a population of uncoupled identical stochastic oscillators that are driven by the same input signal (Figure 1B). This model allows to directly set the number of oscillators in the population (population size, *n*) and the noise intensity of the individual oscillators (*σ*). An intuitive strategy to minimize the stochastic fluctuations at the population level is to increase the number of oscillators in the population (Figure 1C). This result illustrates that even a population recording stems from inherently stochastic oscillators. This stochasticity may influence the population behavior, even if not apparent after averaging over the population of oscillators.

**Figure 1:**
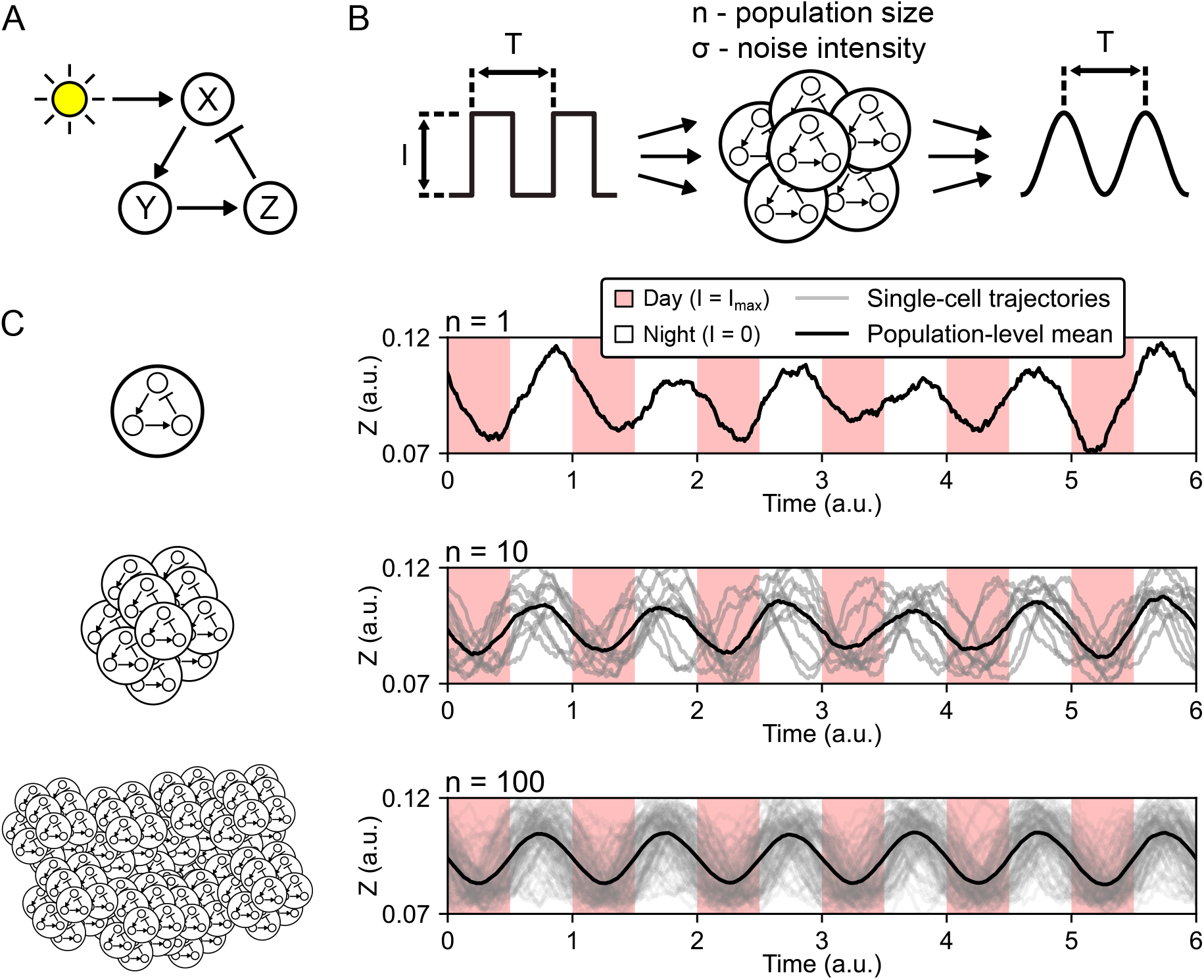
A minimal model of a population of uncoupled cellular oscillators. (A) A minimal oscillator model consists of three variables (*X,, Z*) connected in a negative feedback loop. Input, e.g., light in the case of the circadian clock, is implemented via addition to *X*. (B) The model from panel A represents a single-cell oscillator in a population. A number of oscillators are combined into a population that is driven by a square signal with period *T* and amplitude *I*. The input square signal switches between a high state and a low state, e.g., day and night of the day-night cycle. The output is calculated as the mean of the output of the individual oscillators and, if entrained, oscillates with the same period *T* as the input signal. We aim to explore to parameters of the population of oscillators: the number of oscillators in the population (population size, *n*) and the noise intensity (*σ*). (C) Example results of numerical simulations for a single-cell oscillator (*n* = 1) and two populations of different sizes (*n* = 10, 100). A higher population size results in a smoother periodic signal and thus reduces stochastic fluctuations.

### Noise widens range of entrainment in oscillator populations

Arnold tongues visualize how the entrainment of the oscillator population to the external periodic signal depends on the amplitude and period of the input signal (23, 24). In general, a greater amplitude of the input signal increases the range of periods for which the system entrained, resulting in the typical, tongue-shaped regions of entrainment (Figure 2A). It has been previously shown that if a population mean is considered, the Arnold tongue is wider in comparison to the Arnold tongue estimated with a single stochastic or deterministic model (14). We were interested in how this widening of the entrainment range depends on the noise intensity and the number of oscillators used to construct the population mean. To quantify the Arnold tongues with a single number, we empirically defined an “entrainment area” as the region of the diagram that exhibits entrainment (Figure 2A). Applying this metric to the output of a deterministic model (*σ* = 0) and a stochastic model (*σ* = 0.005), we found that the entrainment area for the deterministic system is lower than for a stochastic population of 1000 oscillators (Figure 2A). This is in line with the previously published findings (14). However, if we consider a stochastic population of only 10 oscillators, the entrainment area drops to a value close to the deterministic model (Figure 2A). This suggests that noise must be compensated for with a sufficiently large population size to allow the wide Arnold tongue.

**Figure 2:**
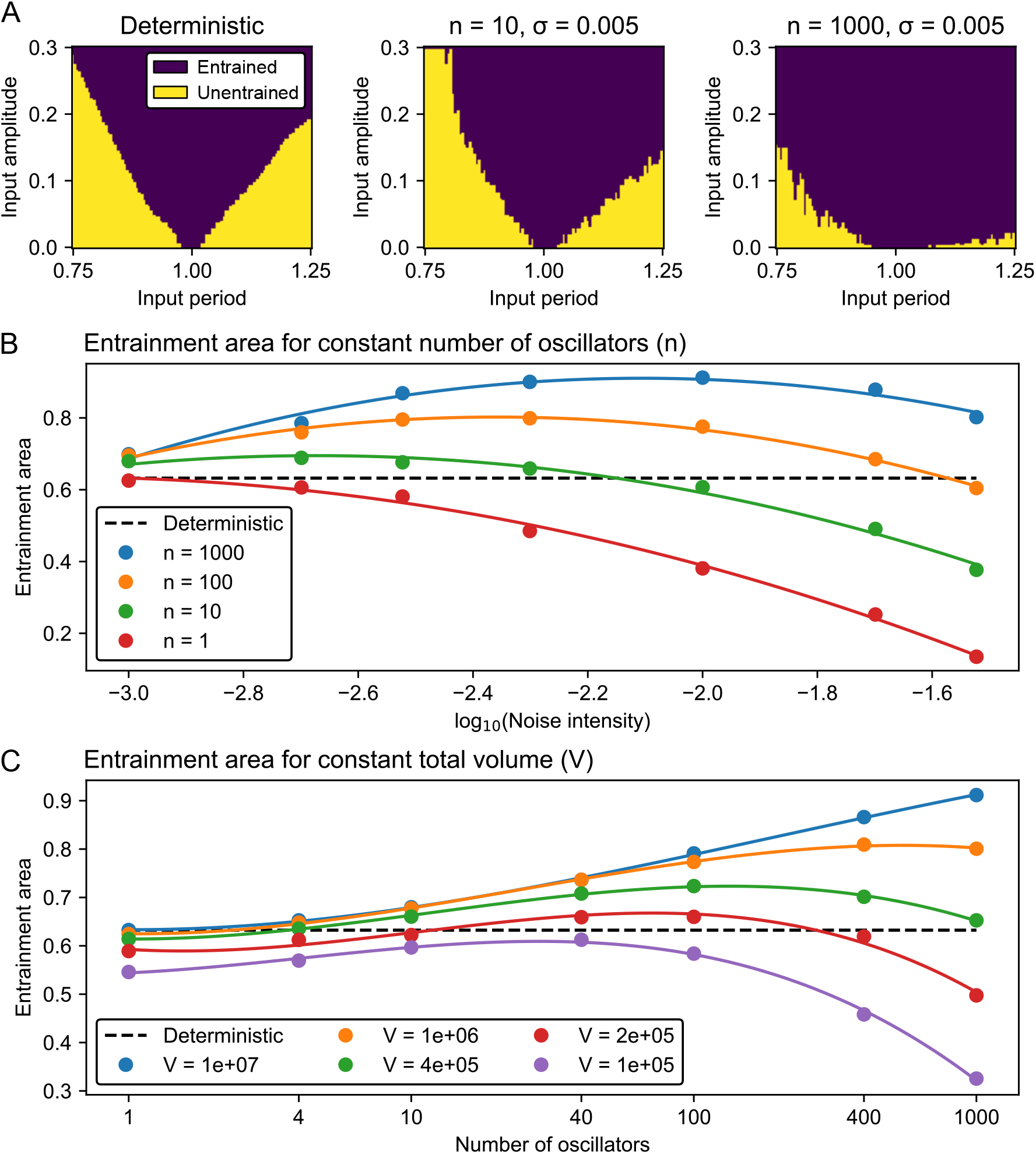
Noise widens the range of entrainment in oscillator populations. (A) Arnold tongue for a deterministic model (*σ* = 0) and a stochastic model (*σ* = 0.005) simulated with two different population sizes. Using an Arnold tongue as a visual cue, we can describe the entrainment area (EA) as the fraction of the diagram where entrainment is observed (blue area, for details see Methods). For the deterministic model, we obtain EA = 0.63. For the stochastic models we obtain EA = 0.66 and EA = 0.90 for the population sizes of *n* = 10 and *n* = 1000 oscillators respectively. (B) Dependence of entrainment area on the noise intensity. The individual Arnold tongues used to generate this image are shown in SI Figure 8. Solid lines are second-order polynomial fits used to estimate the noise intensities for which the entrainment area is maximal (SI Figure 9A). (C) Dependence of the entrainment area on the population size. The individual Arnold tongues used to generate this image are shown in SI Figure 10. Solid lines represent third-order polynomial fits used to estimate the noise intensities for which the entrainment area is maximal (SI Figure 9B).

We extended on this observation by measuring the entrainment area for four different population sizes and seven noise intensities (Figure 2B, SI Figure 8). Using the deterministic case as a reference, we observed that for a single oscillator (*n* = 1) the entrainment area decreases with increasing noise intensity, showing the detrimental effect of noise when only a single oscillator is considered. With increasing population size (*n* > 1), however, an optimal value of noise intensity exists, for which the entrainment area is maximal. With increasing population size, this optimal noise intensity moves to higher values, and also the maximal entrainment area at this optimum noise intensity is increased (SI Figure 9A). Our results suggest an optimal noise intensity for the entrainment of a population of a given size. When this optimum noise intensity is exceeded, the entrainment capacity is again compromised.

To understand how a given population of cellular oscillators (cells) might be tuned towards optimal entrainment, we investigated a hypothetical case where a total volume *V* of cell material is available, but the number of cells (population size, *n*) and their volume (Ω) can change. In other words, we start with one big cell (Ω = *V*) that is subsequently divided into many small cells (Ω = *V*/*n*) while the total volume *V* remains constant. Here, the noise intensity (*σ*) depends on the number of molecules in a cell (Ω) as 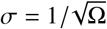 (20, 21). Our simulation results suggests an optimal population size for each total volume (Figure 2C). Specifically, with increasing total volume, the maximal entrainment area occurs at higher population size and the maximal entrainment area itself is also larger (Figure 2C, SI Figures 9B, 10). Meaning, higher total volume can support higher number of noisy oscillators, which in turn allow the population to take advantage of high noise intensities.

### Noise increases phase response to perturbations

We were also interested whether noise can change the responsiveness of the population of uncoupled oscillators to perturbations applied during constant conditions when the input signal is kept zero. This can be explored using phase response curves (PRCs) that plot the change in the oscillator phase caused by a step pulse as a function of the time during the oscillation cycle at which the pulse was applied (25, 26). In circadian research, the PRCs are often characterized based on their amplitude, which represents the extent of the pulse-induced phase shift, as type 1 or type 0. Type 1 PRCs exhibit relatively small phase shifts and appear continuous in the PRC plot, whereas type 0 PRCs show large phase shifts and appear visually discontinuous (25, 26). We have found that the PRC amplitude increases markedly not only with the increasing amplitude of the input signal but also with increasing noise intensity (Figure 3). Specifically for low input amplitude and low noise intensity, we observed relatively small phase shifts (type 1 PRC, Figure 3A, B). As expected, when we increased the input amplitude, the phase shifts also increased. Interestingly, we observed the same effect also by increasing the noise intensity. For high noise intensities, we observed large phase shifts (type 0 PRC) regardless of the amplitude of the input signal (Figure 3C, F, I). Accordingly, increasing noise intensity allows the transition from low-amplitude (type 1) to high-amplitude (type 0) PRC, even if the input amplitude remains low.

**Figure 3:**
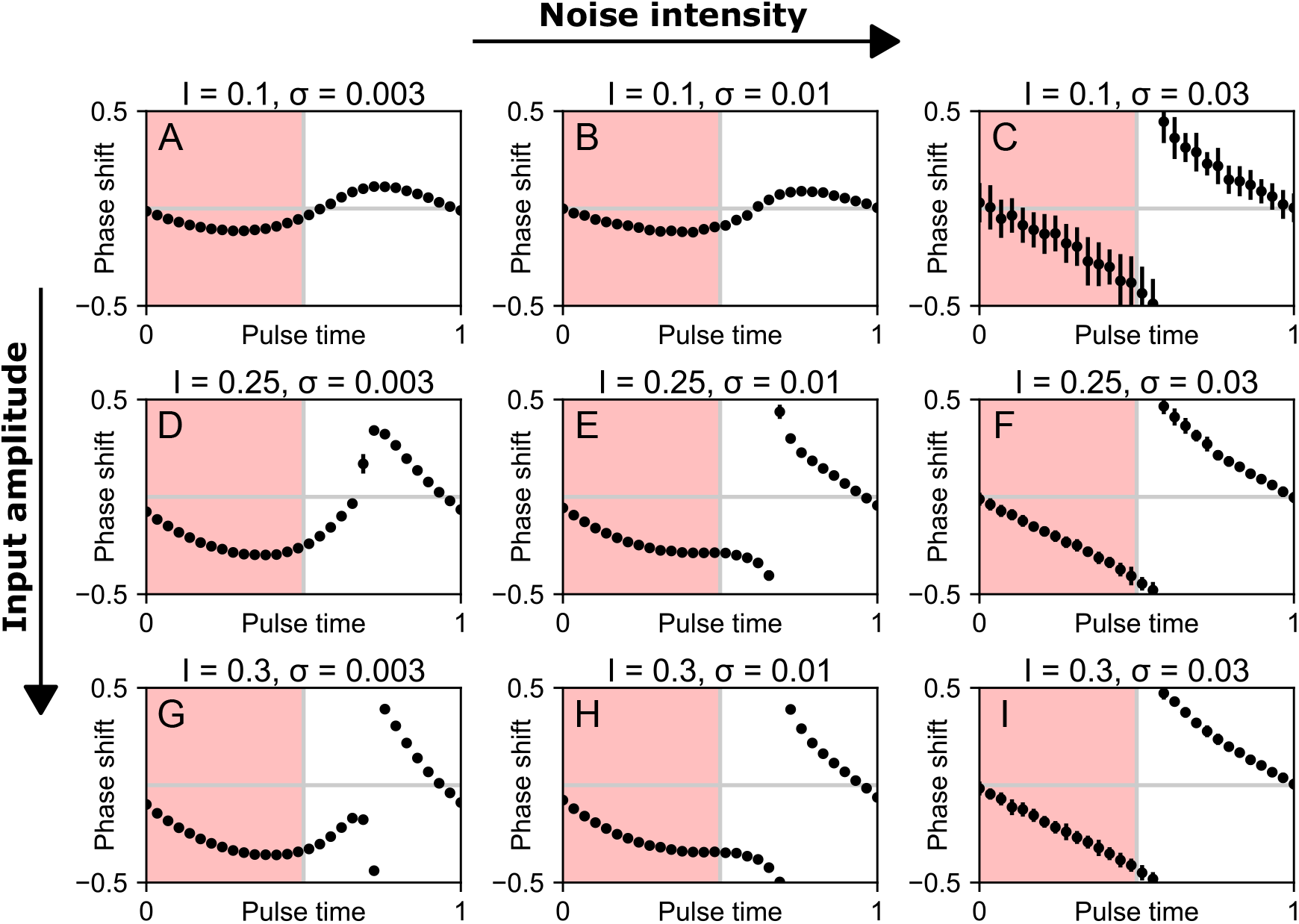
Phase response curves (PRCs) for varying noise intensity (*σ*) and input amplitude (*I*). Phase shift and pulse time are cyclic quantities normalized to the unit circle. The pulse length is 0.5, which is half of the free-running period for the deterministic model (*σ* = 0). The dots represent mean and error bars standard deviation over 10 simulations. (A, B) For low-amplitude input signal and low noise intensity, we obtain a small-amplitude (type 1) PRC. (C) For low-amplitude input signal and high noise intensity, we obtain a large-amplitude (type 0) PRC. (D, E, F, G, H, I) A similar effect of the increasing phase response can be observed also for higher input amplitudes. Note that for high noise intensity (panels C, F, I) we obtain always a large-amplitude (type 0) PRC regardless of the input amplitude.

### Noise allows faster recovery after jet lag

We also explored the effect of a varying noise intensity on the reentrainment to a persistent phase-shift in the input signal, or, “recovery from jet lag” (Figure 4). In these simulations, the population is first entrained by a regular input cycle representing a day-night cycle. After the output of the population is phase-locked to the input signal, an abrupt shift in the phase of the input signal is introduced and time until the population output is locked to the new cycle is measured (SI Figure 11). We found that noise shortens this time, allowing faster recovery from jet lag. This is well visible when the input amplitude is low (Figure 4A). With increasing input amplitude (Figure 4B, C) reentrainment to the new cycle is fast for all noise intensities, so that the effect of the increasing noise is not as apparent. Accordingly, noise allows faster recovery from jet lag, especially if the input amplitude is low.

**Figure 4:**
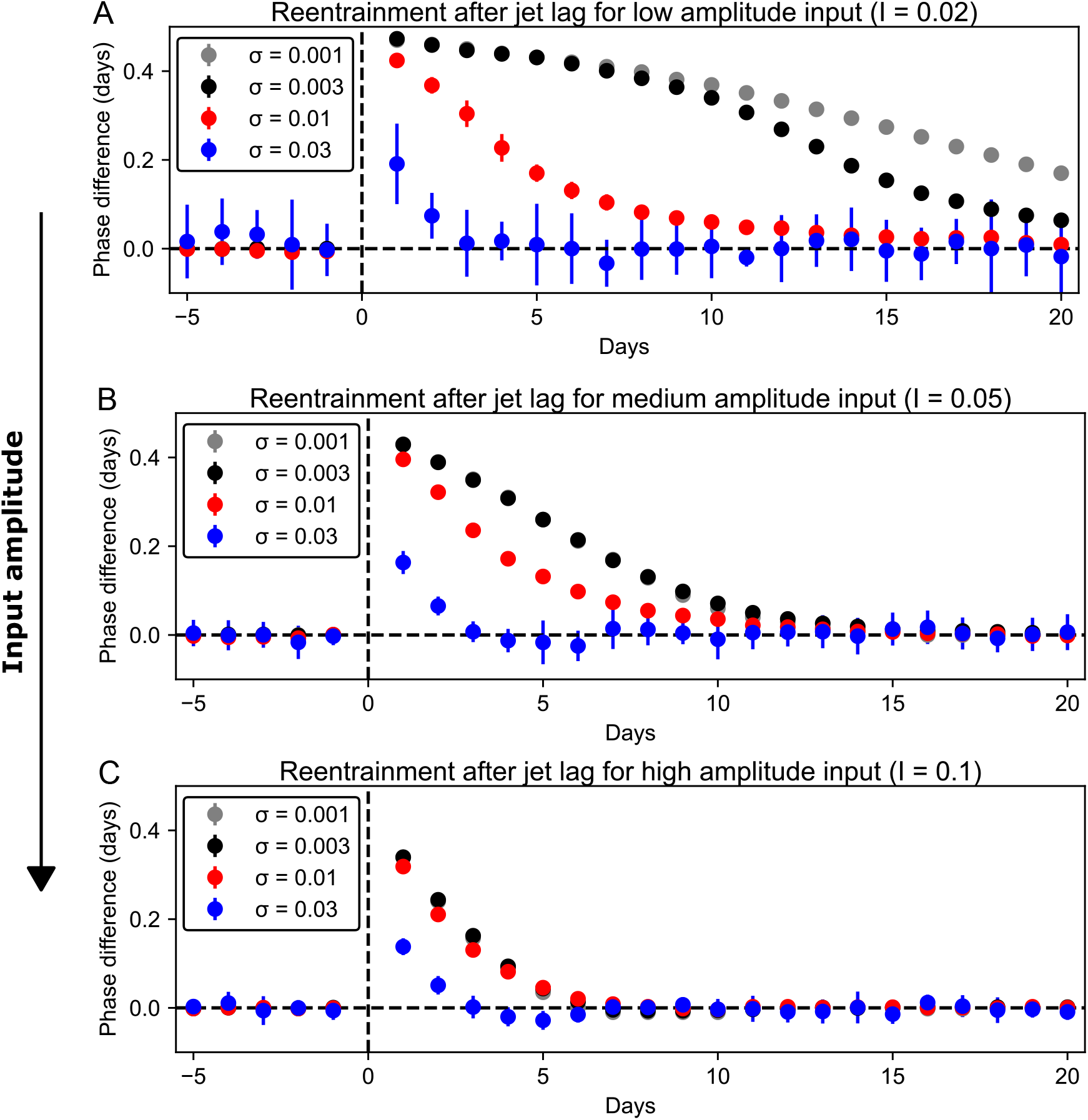
Increasing noise intensity (*σ*) allows faster reentrainment after jet lag, especially for low input amplitudes (*I*). (A) For a low-amplitude input signal, higher noise intensity clearly lead to faster reentrainment. (B) For a medium-amplitude input signal, the difference among the lower noise intensities is shrinking. (C) For a high-amplitude input, the reentrainment after jet lag is fast for all noise intensities.

### Noise facilitates the entrainment of a limit cycle oscillator, but not of a noise-induced oscillator

To verify that the observed phenomena are not only a property of a negative-feedback cellular oscillator but rather a general property of a limit cycle oscillator, we repeated all our experiments with the canonical Van der Pol model (27). The Van der Pol model is a prototypic abstraction for limit cycle oscillators and is used in various fields in science and engineering (28), including biological oscillators such as circadian clock (29) and cardiac pacemakers (30). We used the Van der Pol model with additive input, noise terms, and parameters as used previously to study the entrainment of a stochastic oscillator (22). We explored the behavior of the Van der Pol model under two different parameter sets, one corresponding to a limit cycle oscillator and one corresponding to a noise-induced oscillator (Figure 5A). A noise-induced oscillator is a damped oscillator whose parameters are in the vicinity of the Hopf bifurcation, so that noise can generate sustained oscillatory behavior (31, 32). We found that the entrainment dynamics for the limit cycle model is equivalent to the results obtained with the Kim-Forger model in the previous parts of the manuscript. In particular, with increasing noise intensity and sufficient population size we can achieve enlarged Arnold tongues (Figure 5B, SI Figure 12), increased amplitudes in phase response curves (SI Figure 13), and faster recovery from jet lag (SI Figure 14A). In contrast to the limit cycle oscillator, the noise-induced oscillator showed wide Arnold tongues (Figure 5C, SI Figure 15), high-amplitude PRCs (SI Figure 16), and short jet lags (SI Figure 14B) already for the deterministic model with zero noise intensity. Those properties were consequently not further improved by increasing the noise intensity. We thus conclude that the observations made in this study are general properties of limit cycle oscillators but do not apply to noise-induced oscillators.

**Figure 5:**
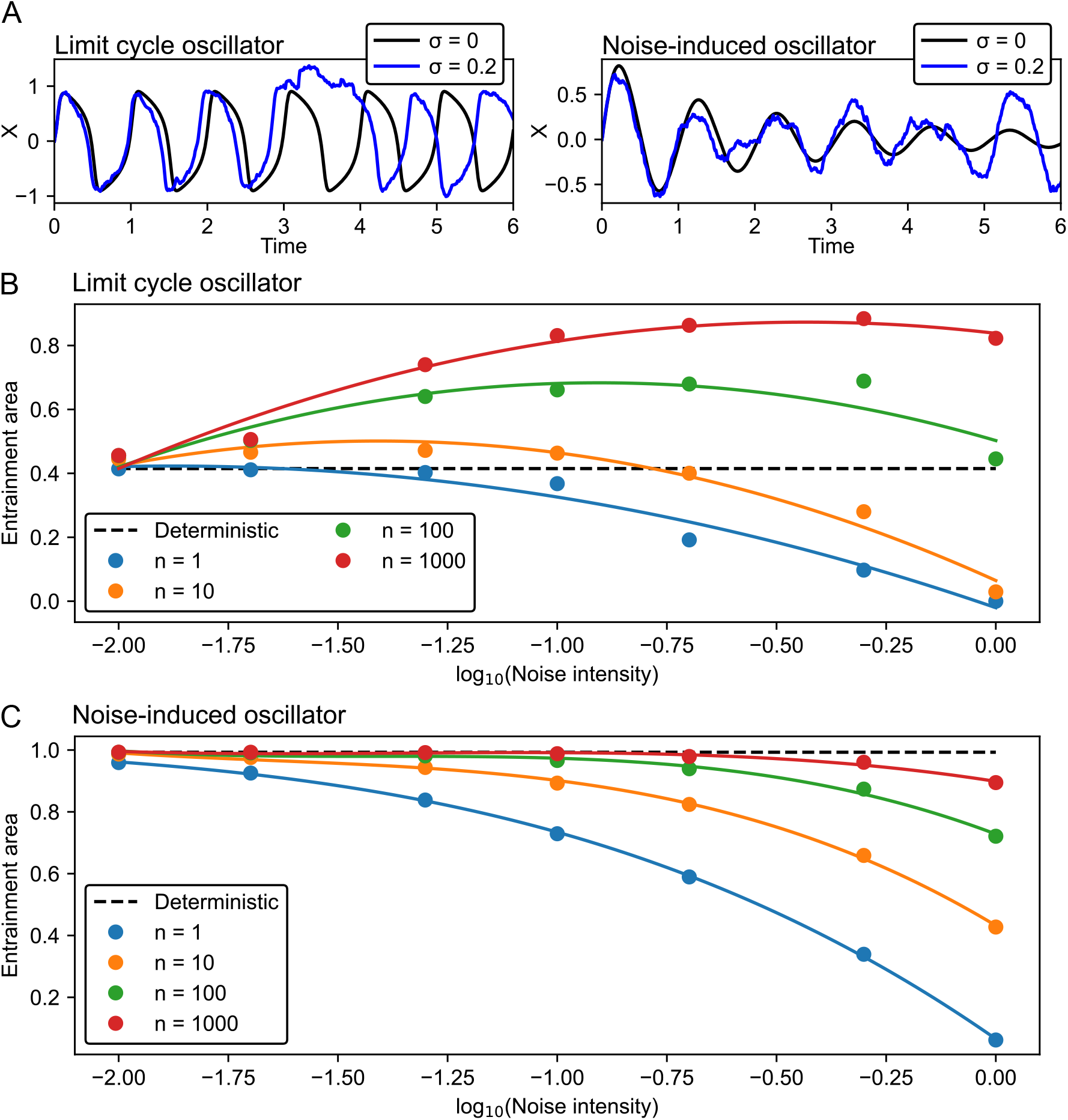
Noise widens the entrainment area for a limit cycle but not for a noise-induced oscillator. (A) Single-oscillator example traces comparing deterministic (*σ* = 0) and stochastic (*σ* = 0.2) simulations for the Van der Pol model with parameters corresponding to a limit cycle and noise-induced oscillators. (B, C) Dependence of entrainment area on the noise intensity for a limit cycle (panel B) and noise-induced (panel C) oscillator. The individual Arnold tongues used to generate those images are shown in SI Figures 12, 15. Solid lines represent second-order (panel B) and third-order (panel C) polynomial fits.

## DISCUSSION

In this work, we explored the entrainment of a population of uncoupled stochastic oscillators represented by a minimal model of the circadian clock. We found that noise allows for population-level entrainment to a wider range of input signal periods and amplitudes. Noise also facilitates a larger response to external stimuli and faster recovery from jet lag. These effects emerge specifically in populations of oscillators, and cannot be observed in single oscillators. We used the canonical Van der Pol model to show that this enhanced entrainment emerges also for a generic limit cycle oscillator, but not for a noise-induced oscillator. In the field of cellular oscillators and especially circadian clocks, these findings should contribute to a better understanding of the population behavior of cells, for example in cell cultures, and how the population behavior relates to the behavior of single cells. Since our results seem to apply to various limit cycle oscillator systems, the results are not limited to cellular oscillators but can be applied to any domain where a population of noisy oscillators is under periodic pacing.

Previously published results have shown that noise widens the population-level range of entrainment (14). It has been, however, also shown that, in the case of a single stochastic oscillator, the range of entrainment decreases with increasing noise intensity (33). We have extended those findings by exploring the entrainment for a number of population sizes and noise intensities and found that for each population size, an optimal noise intensity exists. This existence of an optimal noise intensity is a typical characteristic of a phenomenon known as stochastic resonance (34). The term stochastic resonance is traditionally used in neuroscience to describe improved detection of weak signals in threshold-like systems (35). The term stochastic resonance is also used more generally to describe improvement in output performance of a noisy system in various disciplines including cell biology, ecology, and physics (36). In our system, noise improves the population-level entrainment but only to the point the population size is sufficiently large to compensate for the noise-induced fluctuations at the population-level read-out. Thus, in the comparison to the previous studies, we showed not only that noise widens the range of entrainment, but also that there exists an optimal value of the noise intensity, for which the range of entrainment is the widest.

In our model of circadian oscillations, the noise intensity (*σ*) and the number of interacting molecules (Ω) are inversely related: lower Ω gives higher *σ* and vice versa (20, 21). We took advantage of this physical interpretation and perform a series of experiments, where the total number of molecules (*V*) is fixed and is divided equally among *n* cells. We found that there is an optimal number of cells, for which the range of entrainment is the widest. In a general system, we could interpret *V* as a total amount of available resources or as a total price of a system and Ω as the amount of resources taken by a single unit. Consequently, we showed that, if the goal is maximal sensitivity in entrainment to input signals, it is more advantageous to distribute the resources to several noisy units rather than maintain one with minimal noise. However, when the population size is increased beyond its optimum, the individual units become too noisy and the ability of entrainment is again compromised.

In this work, we showed that the amplitude of the phase response curves and speed of recovery from jet lag depends not only on the amplitude of the input signal, which might be obvious (37), but also on the noise intensity. In our experiments, higher noise intensities led to high PRC amplitude and shorter jet lags. These results might be particularly interesting in the context of the entrainment dynamics of the circadian clock. In mammals, the clock is thought to be organized hierarchically, with a central pacemaker of highly coupled oscillators in the suprachiasmatic nucleus (SCN) and less coupled peripheral oscillators that are entrained by signals from the SCN (9). Pharmacologically increased noise in the SCN cells shortens jet lag even without explicitly weakening coupling among the cells (38). This hints at the possible applicability of our results also in the domain of coupled oscillators. The coupling also provides the SCN with a certain level of resistance to noise and external perturbation (10, 11). Our results show that for increasing noise intensities the population becomes very sensitive to the input signals. This property would not seem useful to the SCN in maintaining a steady rhythm, but might in fact be advantageous for peripheral clocks that need to adjust to a manifold of external cues.

The positive effect of noise for the circadian clock is usually illustrated by noise-induced oscillations in the vicinity of Hopf bifurcation (31, 32). From circadian data, however, it is often impossible to infer whether the circadian rhythms are formed by limit cycle or noise-induced oscillators (39). It has been shown that a noise-induced oscillator can be entrained to a wider range of input amplitudes than a limit cycle oscillator (40). We studied a general Van der Pol model with additive light and noise terms and with two parameter sets representing limit cycle and noise-induced oscillations (22). Our results showed that indeed noise-induced oscillators entrain easier than limit cycle oscillators not only at the single-cell level but also at the population level. However, the speed and range of entrainment for the limit cycle oscillator improves with increasing noise, whereas the entrainment properties of the noise-induced oscillator remain unchanged. For high noise intensities and sufficient population sizes, the entrainment dynamics of the limit cycle and noise-induced oscillators in fact become increasingly similar in terms of wide range of entrainment, high-amplitude phase response curves, and short jet lags.

Another important application of our results is in inferring the single-cell dynamics from the population-level bioluminescence assays. These assays are relatively cheap and fast to conduct and lend themselves well to high-throughput screening of chemical compounds and mutants (41). It has been shown that the cells in the bioluminescence assays behave as uncoupled oscillators (12, 13). Bioluminescence assays record population mean over thousands of cells so caution is required if we desire to make conclusions about single-cell behavior. A common example of a discrepancy between the population-level and single-cell behavior is the desynchronization of the cells in constant darkness that lead to the absence of oscillations at the population level even though single cell oscillations persist (42). In the context of circadian entrainment, assays based on zebrafish cell lines are especially suitable because zebrafish cells are directly light-responsive (43). These assays were also previously modeled as uncoupled oscillators, reproducing even detailed aspects of the experimental data (17). Considering that noise can be pharmacologically enhanced in cell cultures (44), zebrafish assays might be a suitable experimental system to investigate some of the presented results in further in vitro experiments.

## AUTHOR CONTRIBUTIONS

V.K. carried out the simulations and analyzed the results. V.K., L.H., and R.M. conceptualized the study and wrote the article.

## ACKNOWLEDGEMENTS

We are grateful for funding by the Helmholtz Association in the program Natural, Artificial and Cognitive Information Processing (NACIP), and HIDSS4Health – the Helmholtz Information & Data Science School for Health. We acknowledge support by the KIT-Publication Fund of the Karlsruhe Institute of Technology.

