## Supplementary Information for "Noise facilitates entrainment of a population of uncoupled limit cycle oscillators"

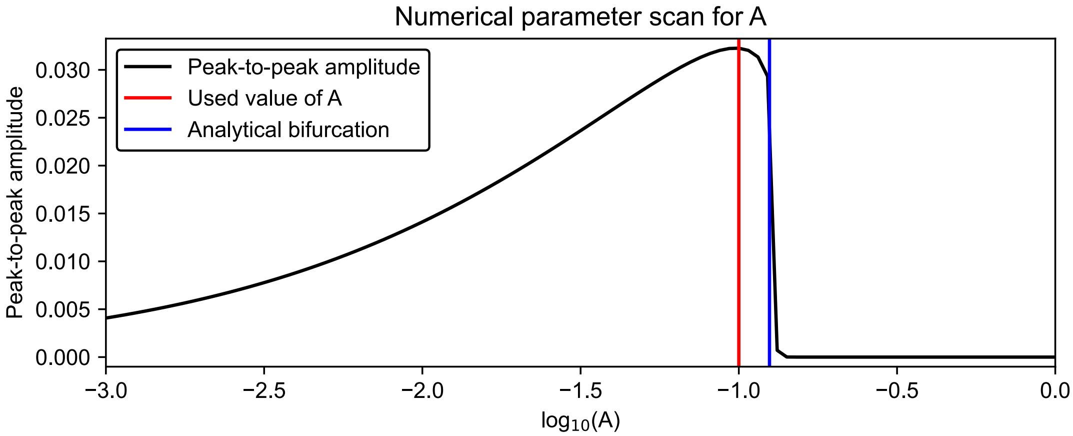


**Figure 1.** Peak-to-peak amplitude of the model variable $Z$ for varying parameter $A$. We set $A = 0.1$ for all numerical experiments in this study. This corresponds to the region with the highest amplitude limit cycle oscillations.

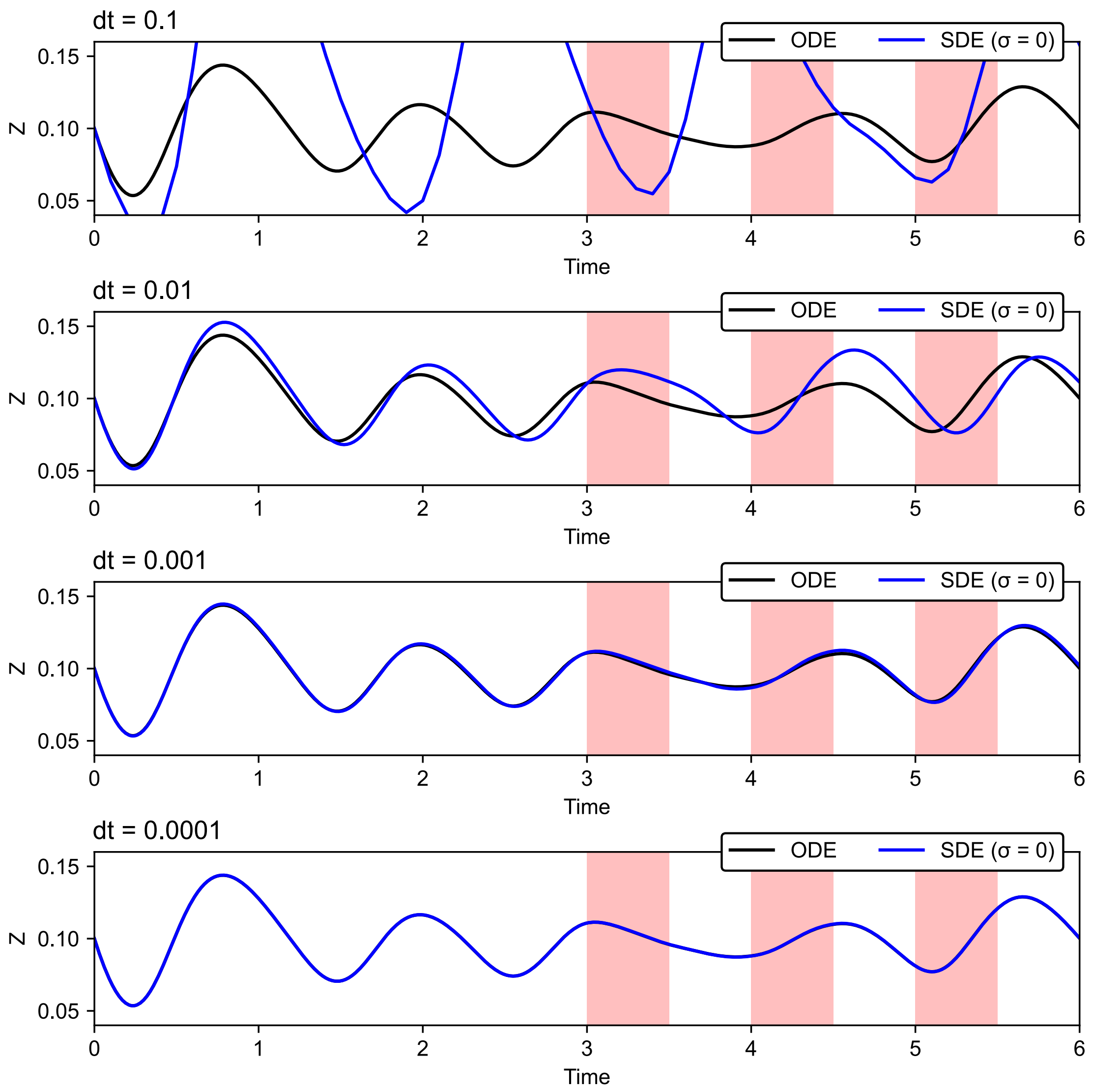


**Figure 2.** The stochastic Kim-Forger model was simulated using the Euler-Maruyama method with a fixed integration step set to dt = 0.0001 (last panel). For $\sigma = 0$, the stochastic numerical simulation gives the same output as an adaptive ODE solver.

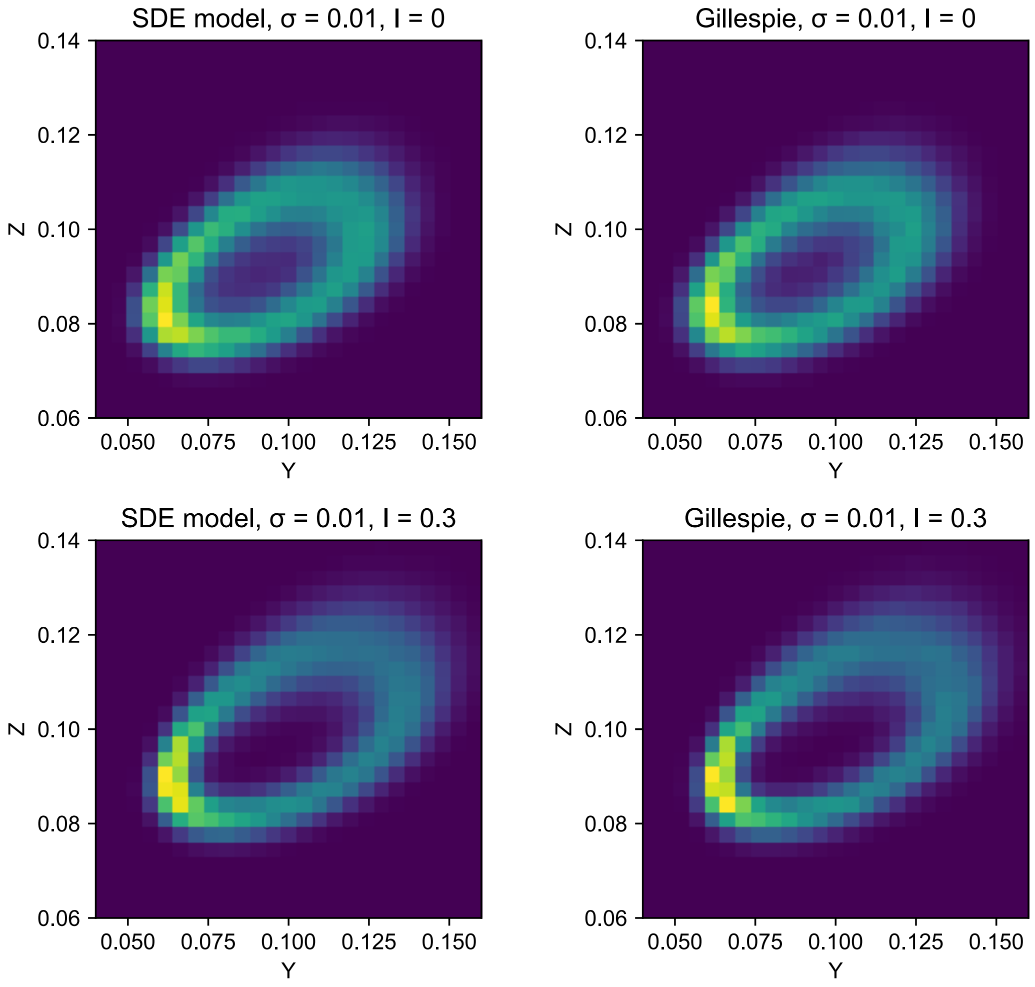


**Figure 3.** Phase space histogram comparing the numerical simulation of the SDE model (first column) and the Gillespie method (second column). The SDE model gives comparable results to the Gillespie method under free running conditions (first row) as well as periodic input (second row). The color code indicates the probability of the model output to visit the specific state with bright yellow indicating high probability and dark blue indicating low probability.

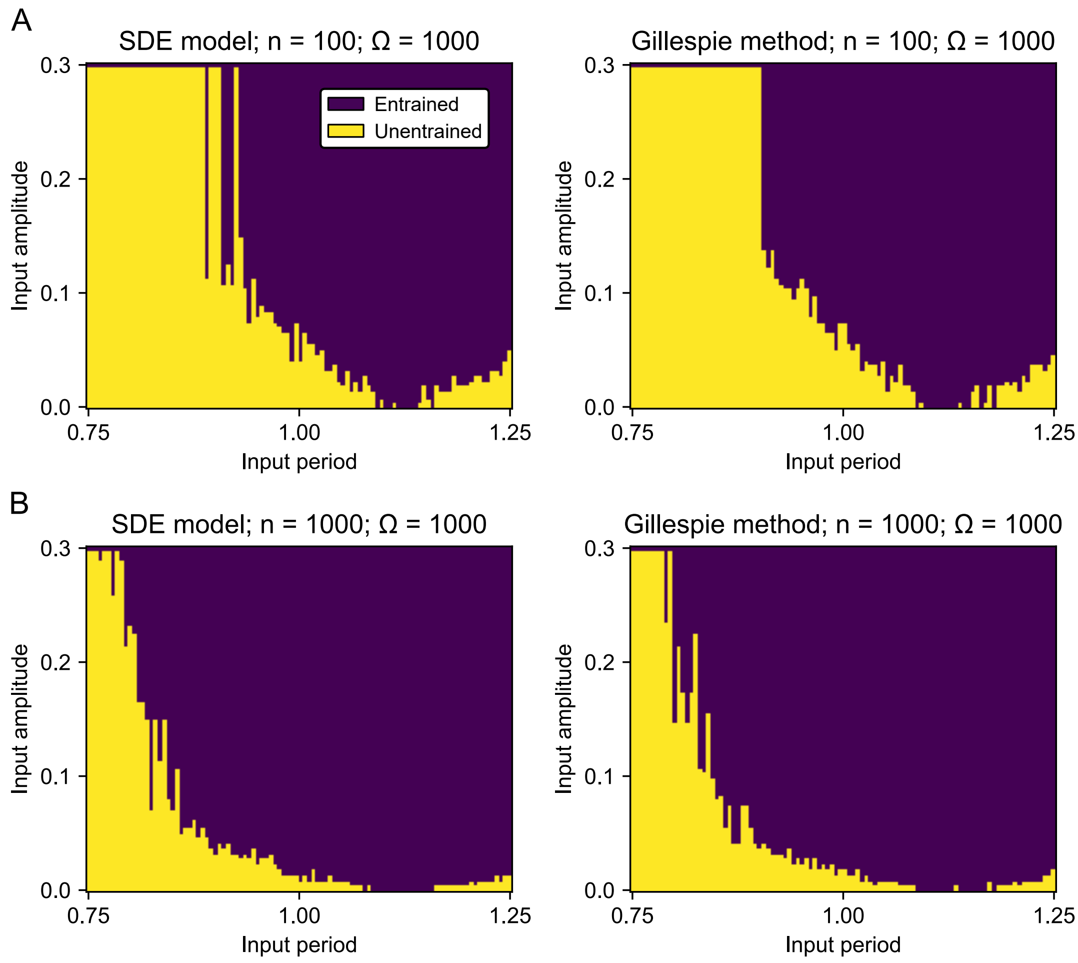


**Figure 4.** For $\Omega = 1000$ the Arnold tongues estimated with the SDE model and Gillespie method are equivalent.

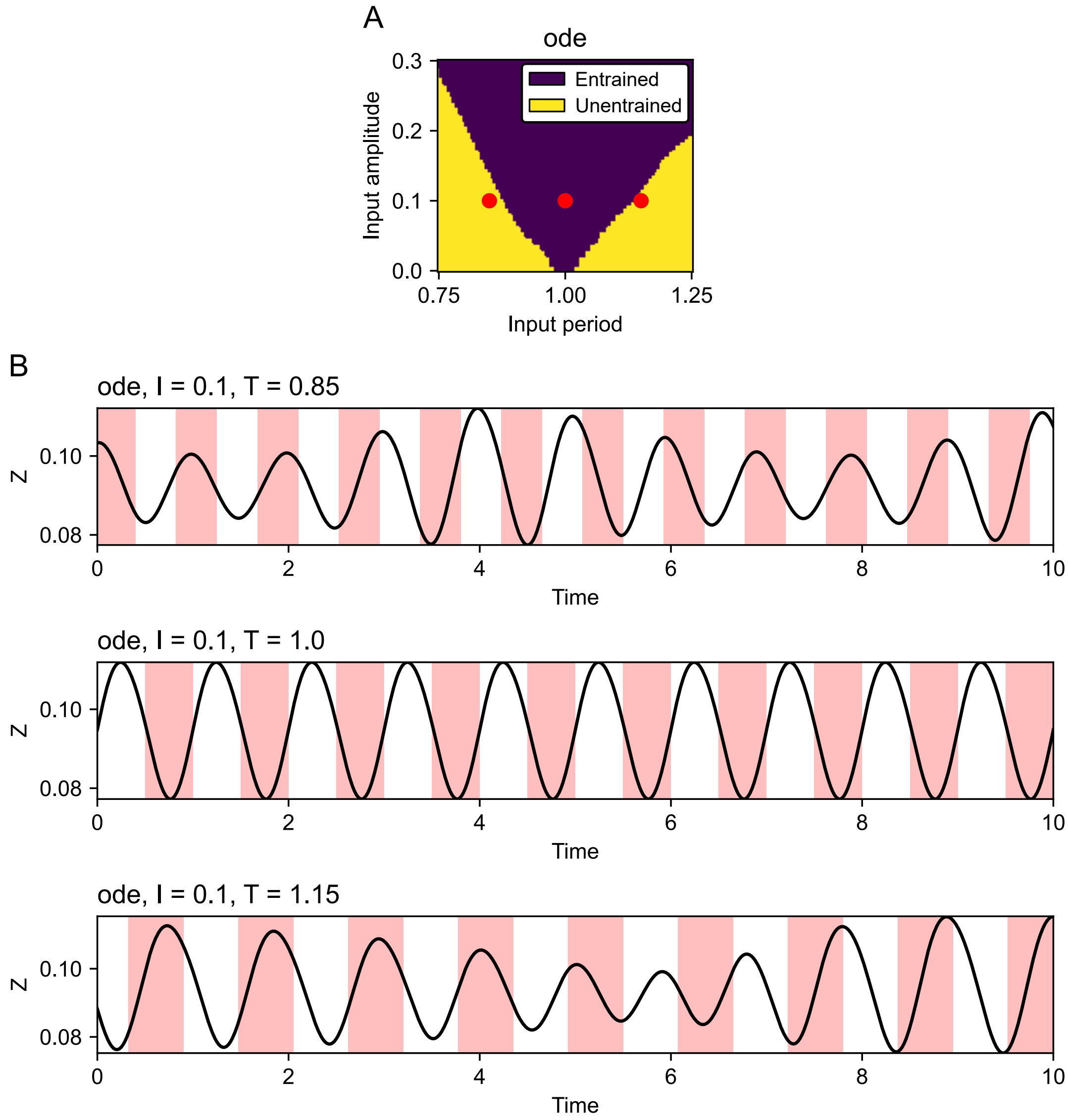


**Figure 5.** Example traces for Arnold tongue generating using a deterministic model. (A) Arnold tongue for the ODE model ($\sigma = 0$). The red dots indicate, from left to right, the time traces from panel B, from top to bottom. (B) Time traces for the same input amplitude and three input periods as indicated by the red dots in panel A. The time series for $T = 1$ is entrained and thus lies in the blue area of the Arnold tongue. The time series for $T = 0.85$ and $T = 1.15$ are not entrained and thus lie in the yellow area of the Arnold tongue. For the definition of entrainment and its estimation see Methods in the main text.

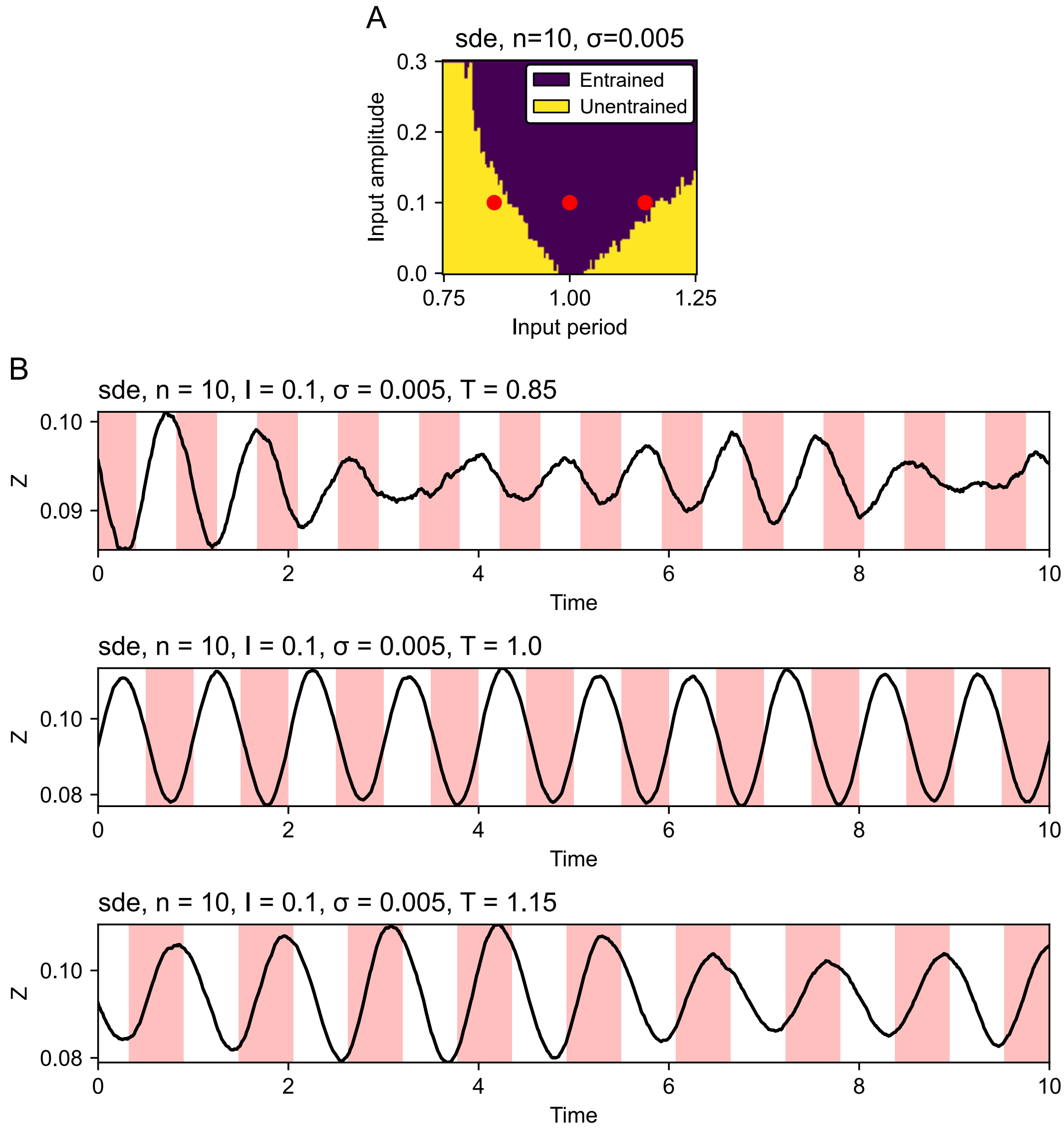


**Figure 6.** Example traces for Arnold tongue for a population of 10 oscillators. (A) Arnold tongue for an SDE model. The red dots indicate, from left to right, the time traces from panel B, from top to bottom. (B) Time traces for the same input amplitude and three input periods as indicated by the red dots in panel A. The time series for $T = 1$ and $T = 1.15$ are entrained and thus lie in the blue area of the Arnold tongue. The time series for $T = 0.85$ is not entrained and thus lies in the yellow area of the Arnold tongue. For the definition of entrainment and its estimation see Methods in the main text.

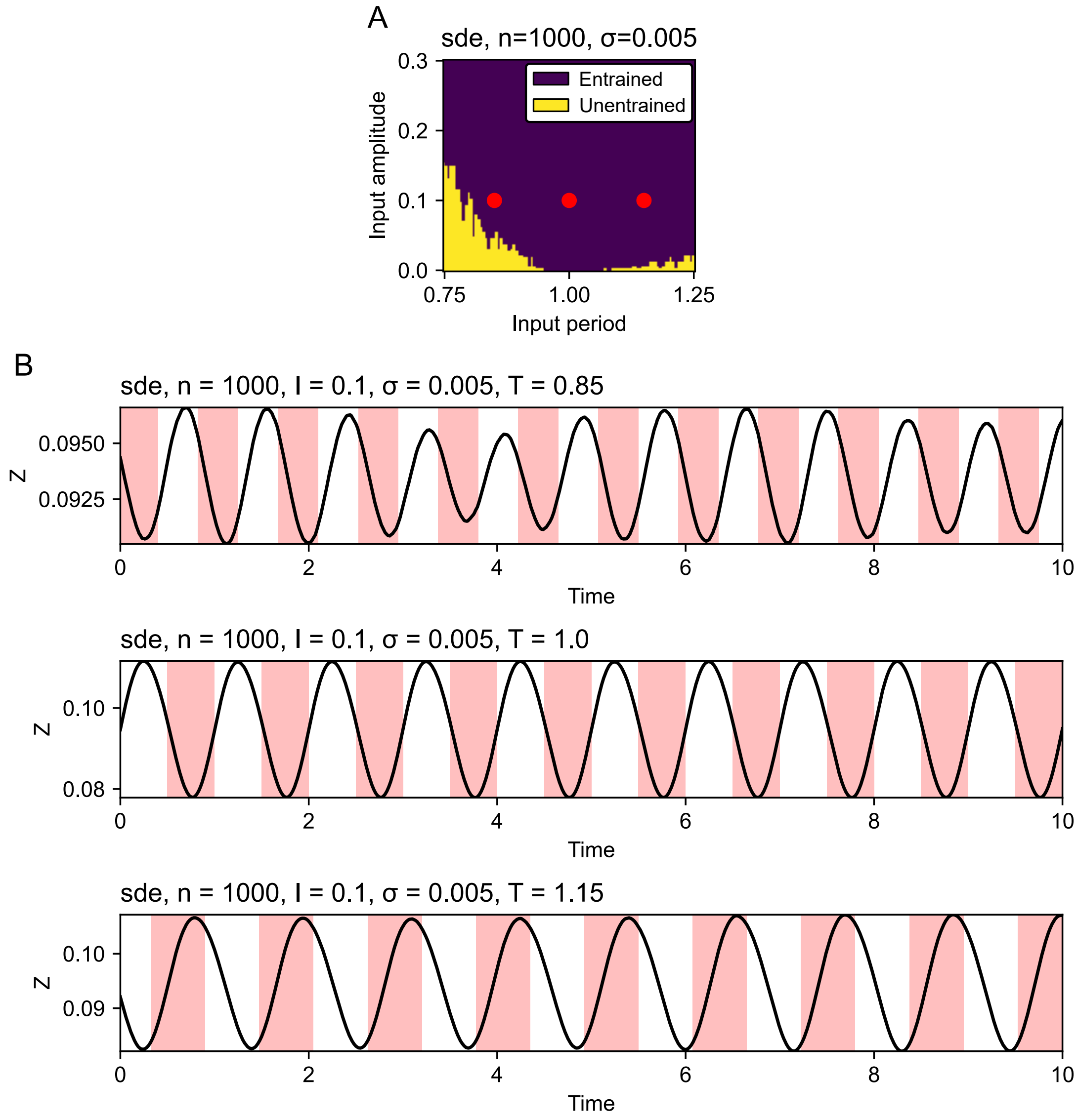


**Figure 7.** Example traces for Arnold tongue for a population of 1000 oscillators. (A) Arnold tongue for an SDE model. The red dots indicate, from left to right, the time traces from panel B, from top to bottom. (B) All three time series are entrained and thus lie in the blue area of the Arnold tongue. For the definition of entrainment and its estimation see Methods in the main text.

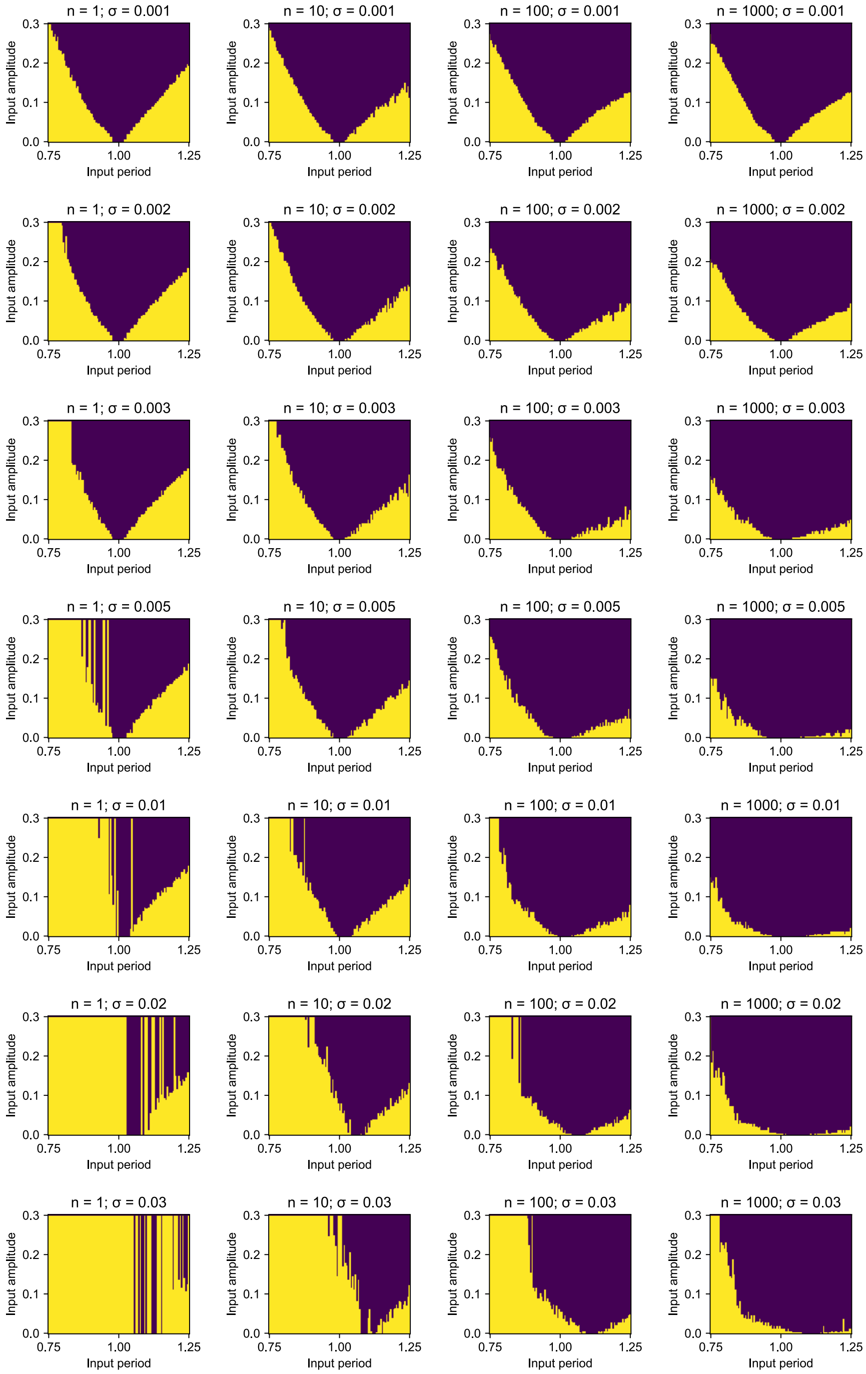


**Figure 8.** Arnold tongues used to calculate the entrainment area for Figure 2B.

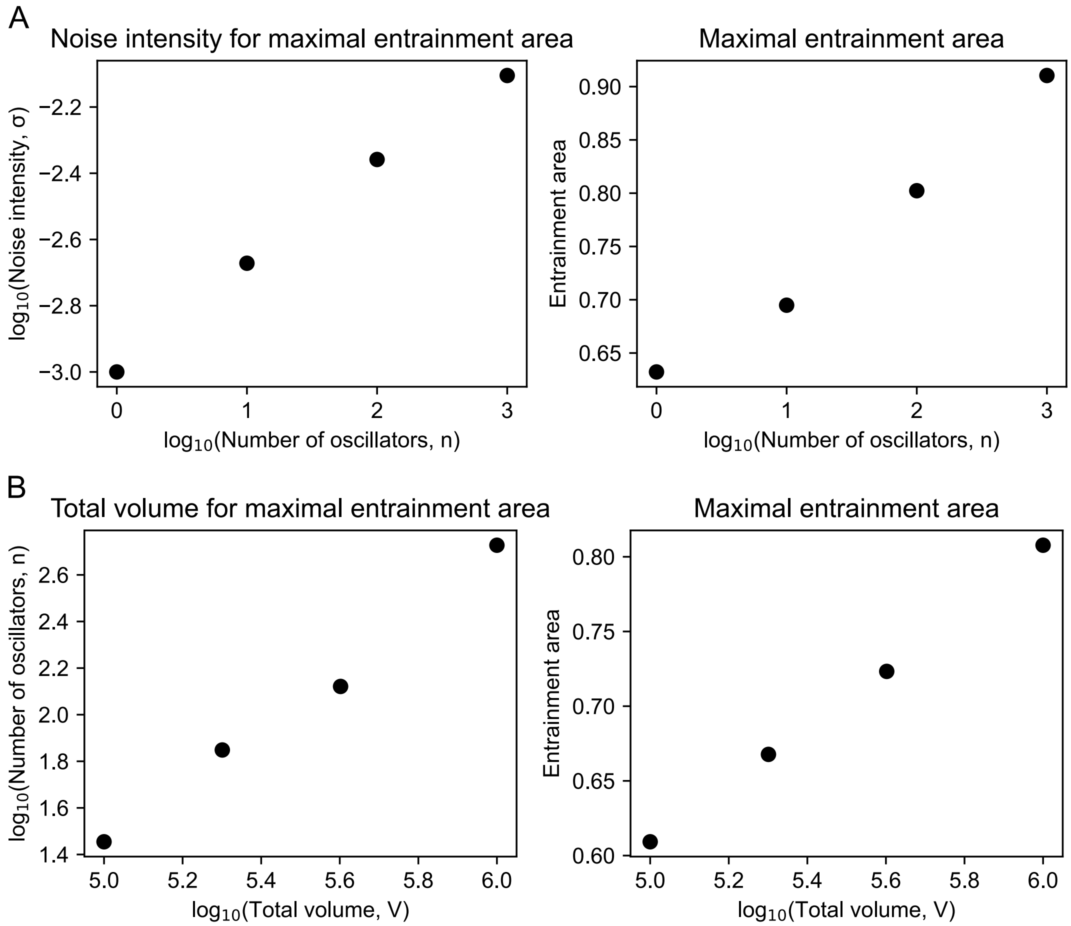


**Figure 9.** A larger population size compensates for a high noise intensity and allows the positive effect of noise to be more prominent. (A) Estimated peak values from Figure 2B. With the increasing total volume, the population size (number of oscillators) also increases, at which the maximal entrainment area occurs (left panel). The maximal entrainment area itself is also increasing with increasing population size (right panel). (B) Estimated peak values from Figure 2C. With the increasing total volume, the population size (number of oscillators) also increases, at which the maximal entrainment area occurs (left panel). The maximal entrainment area itself is also increasing with increasing population size (right panel).

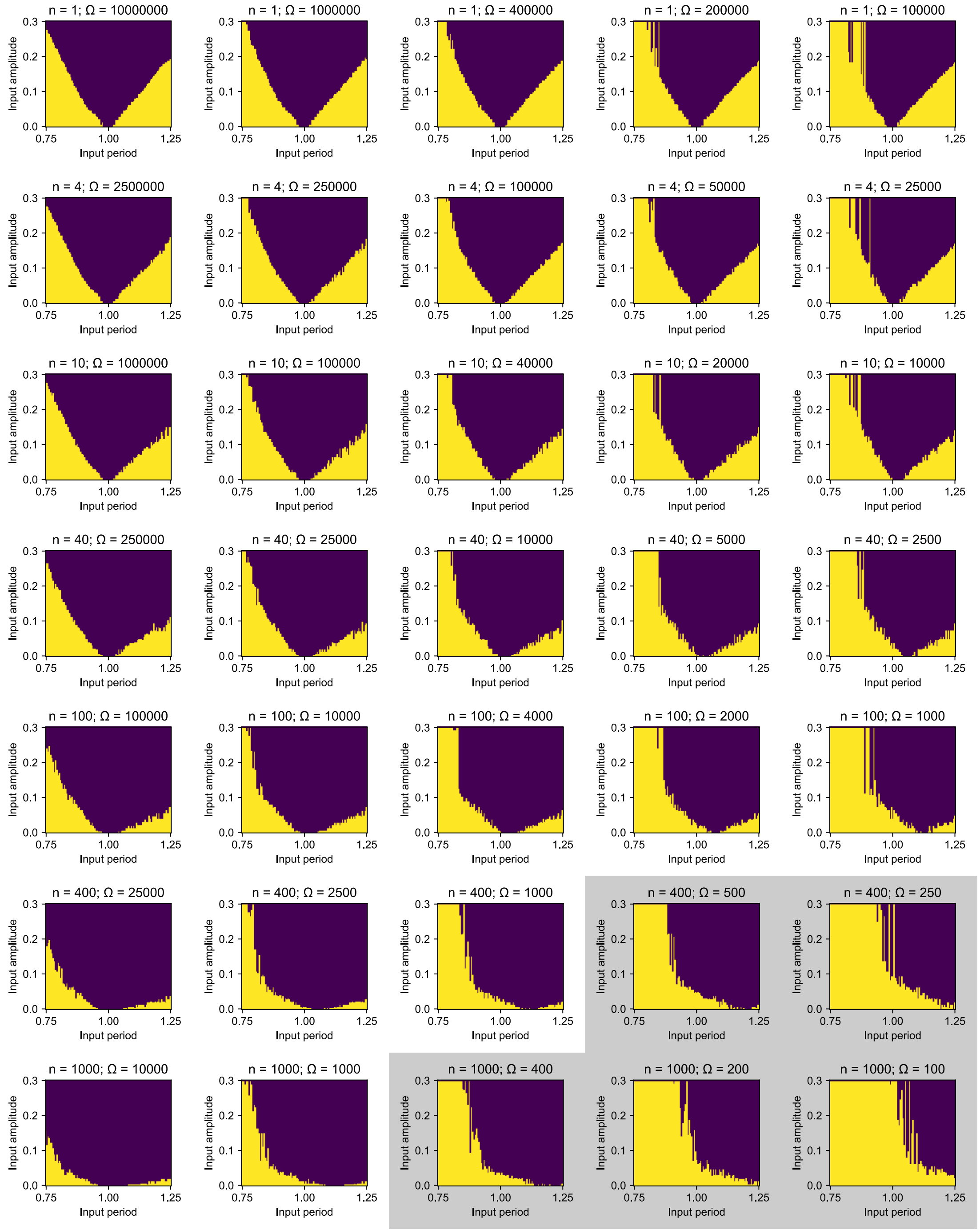


**Figure 10.** Arnold tongues used to calculate the entrainment area for Figure 2C. In gray, for low values of Ω we used Gillespie method instead of the SDE model as described in Methods in the main text.

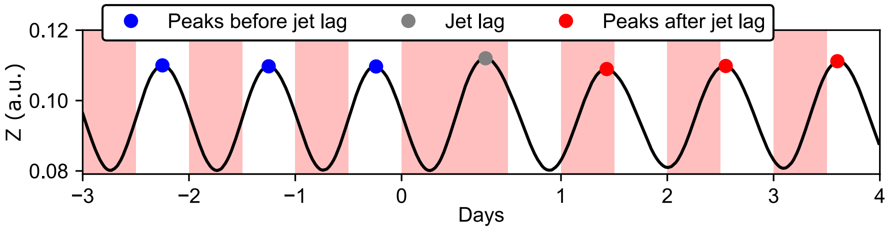


**Figure 11.** Jet lag is simulated by keeping the input square signal in its high state for doubled time and then proceeding again with a regular rhythm.

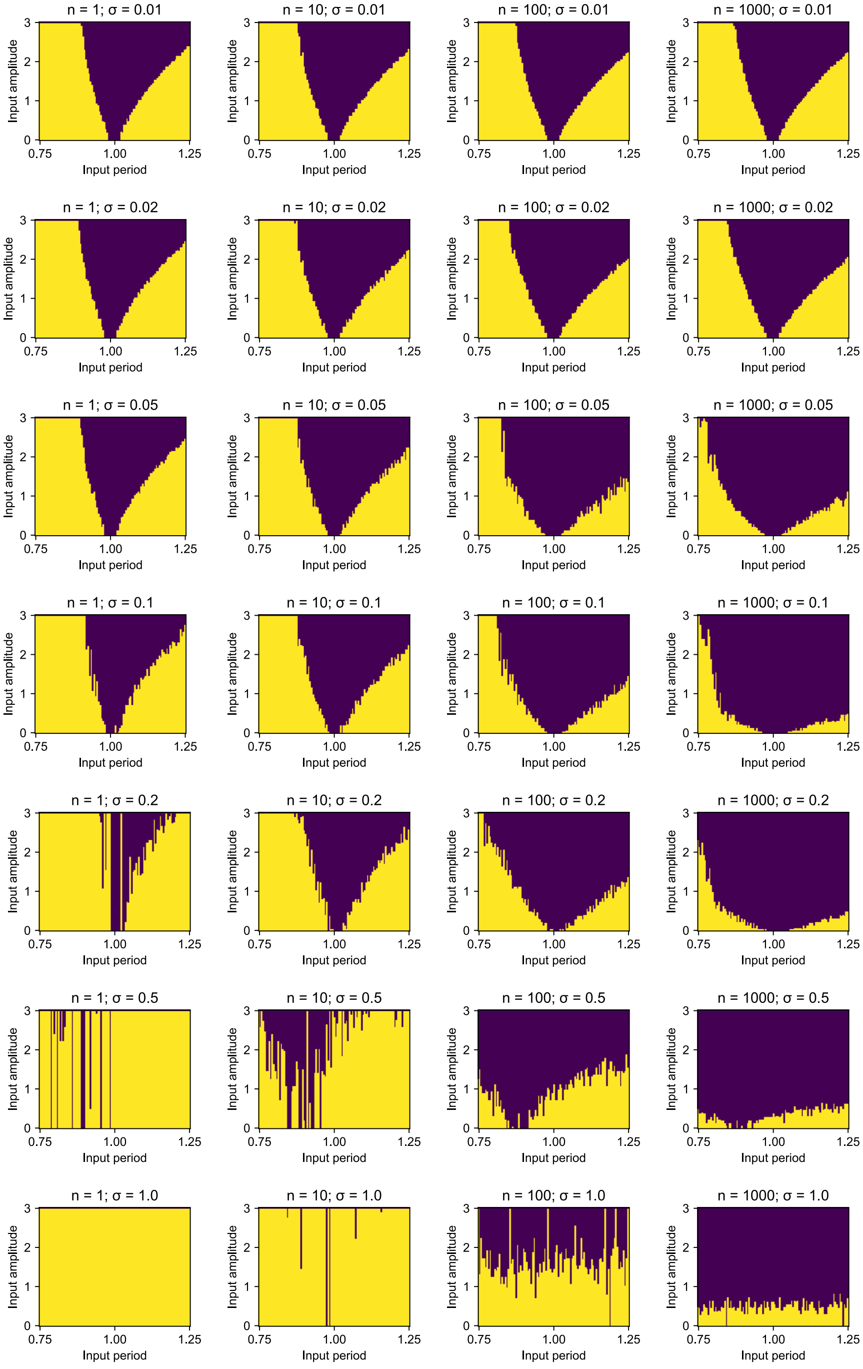


**Figure 12.** Individual Arnold tongues used to estimate the entrainment area in Figure 5B (limit cycle Van der Pol).

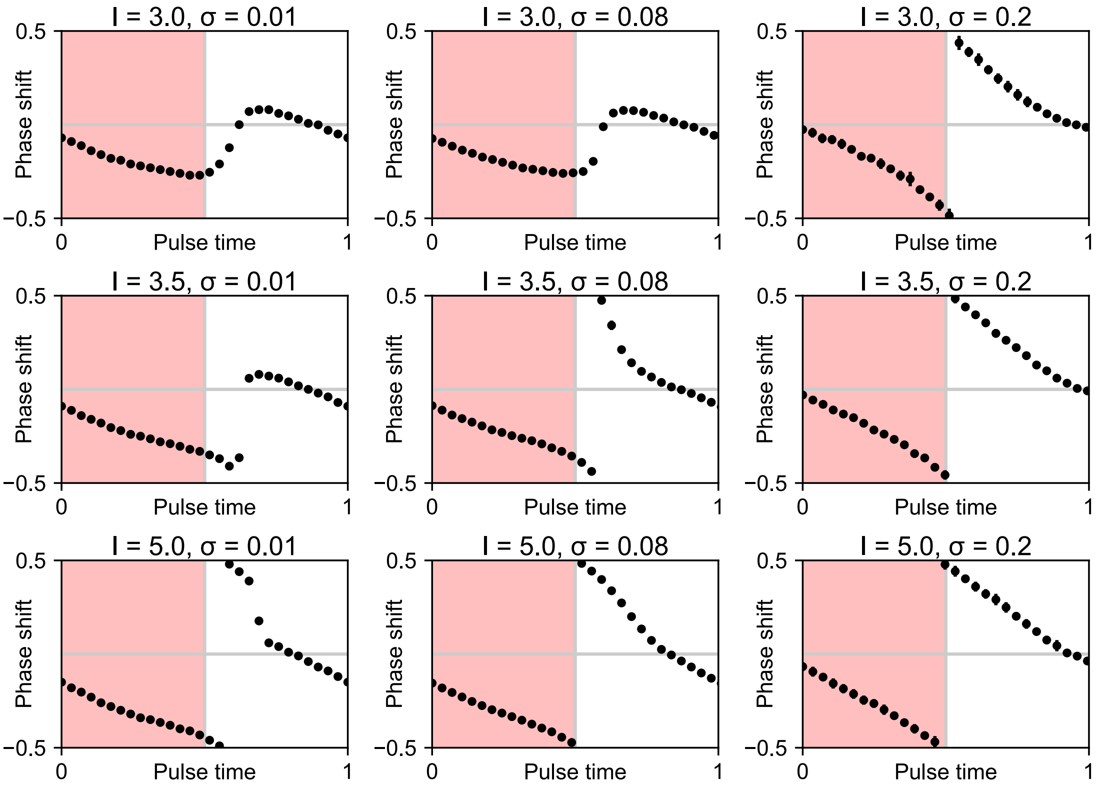


**Figure 13.** Phase response curves (PRCs) for varying noise intensity ($\sigma$) and input amplitude ($I$) for the limit cycle Van der Pol model. Phase shift and pulse time are cyclic quantities normalized to the unit circle. The pulse length is 0.5, which is a half of the free-running period for the deterministic model ($\sigma = 0$). For low-amplitude input signal and low noise intensity (left top) we obtain a small-amplitude PRC. The amplitude of the PRC increases by increasing input amplitude (moving down) or by increasing noise intensity (moving right). Notice that for high noise intensity (right column) we obtain a high-amplitude PRC also for a low input amplitude.

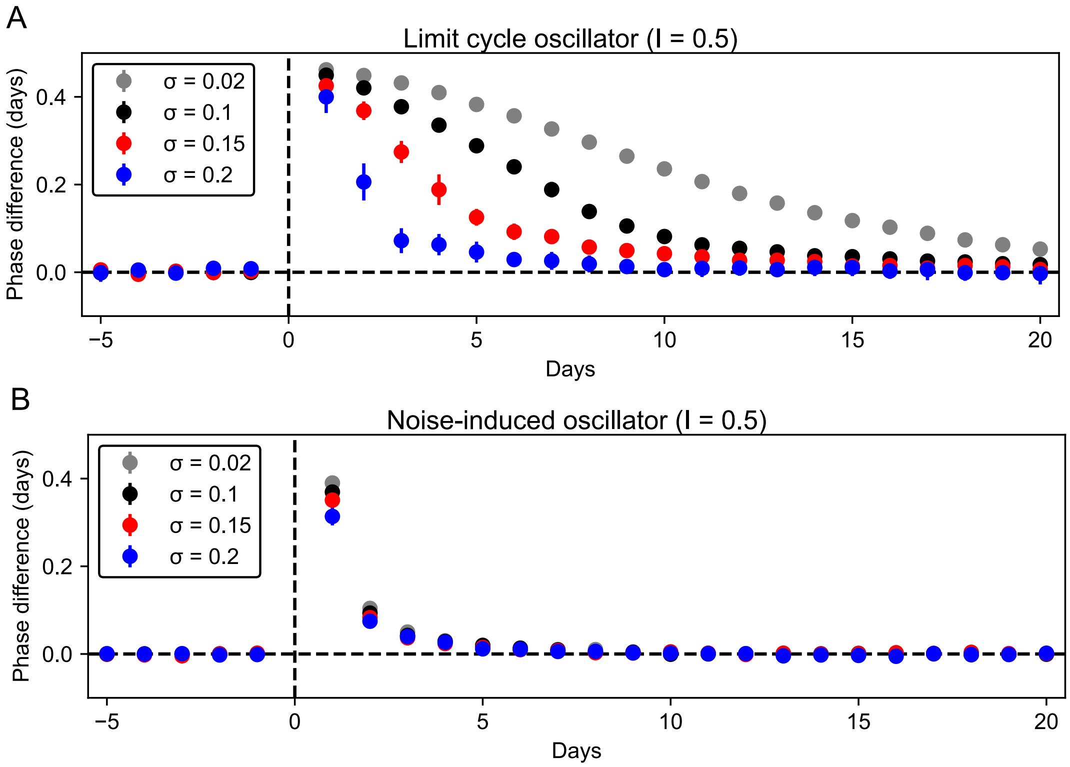


**Figure 14.** Increasing noise intensity ($\sigma$) allows faster reentrainment after jet lag for a limit cycle but not for a noise-induced Van der Pol model. (A) The limit cycle Van der Pol model shows the same behavior as the Kim-Forger model with shortening jet lag with increasing noise intensity. (B) The noise-induced Van der Pol model shows short jet lags regardless the noise value.

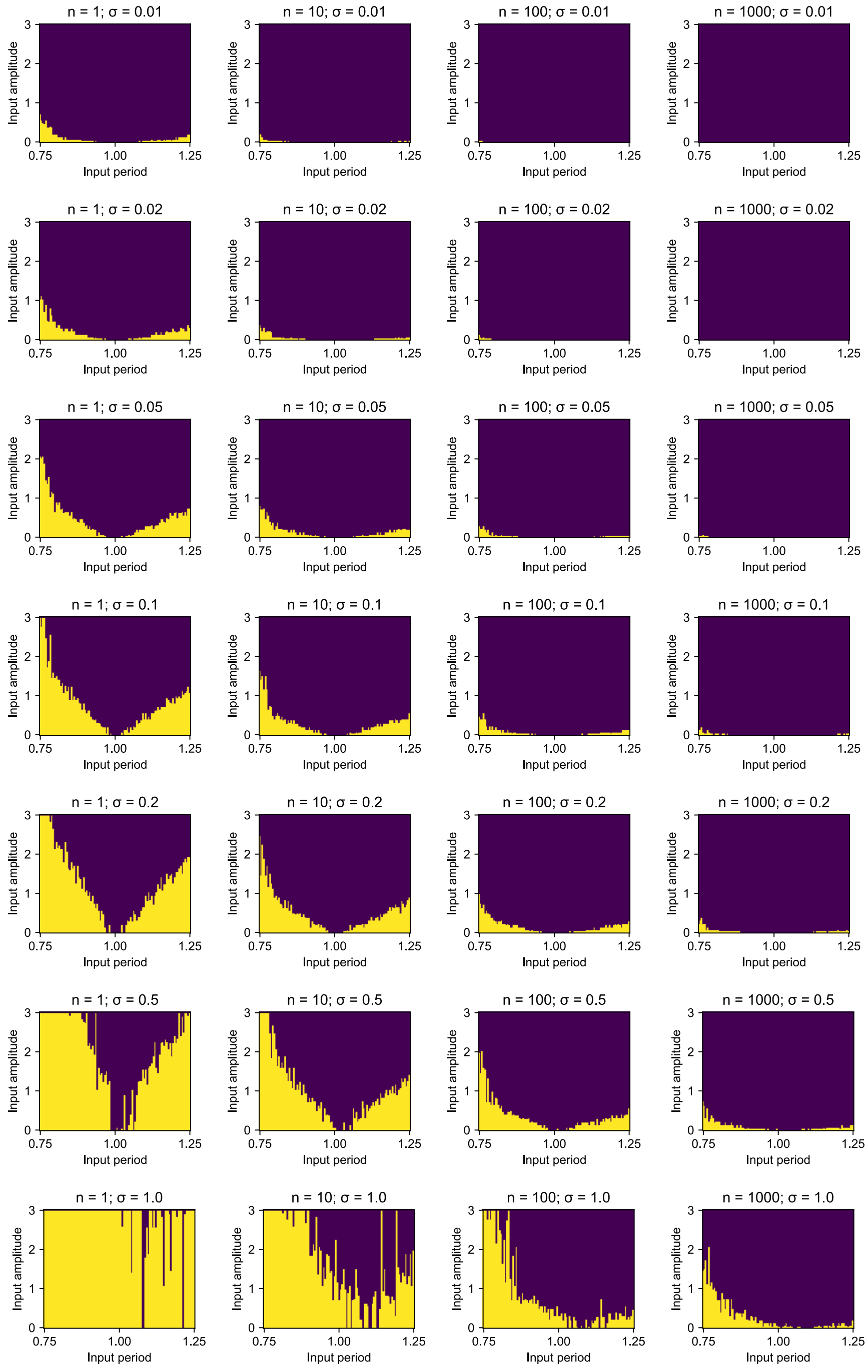


**Figure 15.** Individual Arnold tongues used to estimate the entrainment area in Figure 5C (noise-induced Van der Pol).

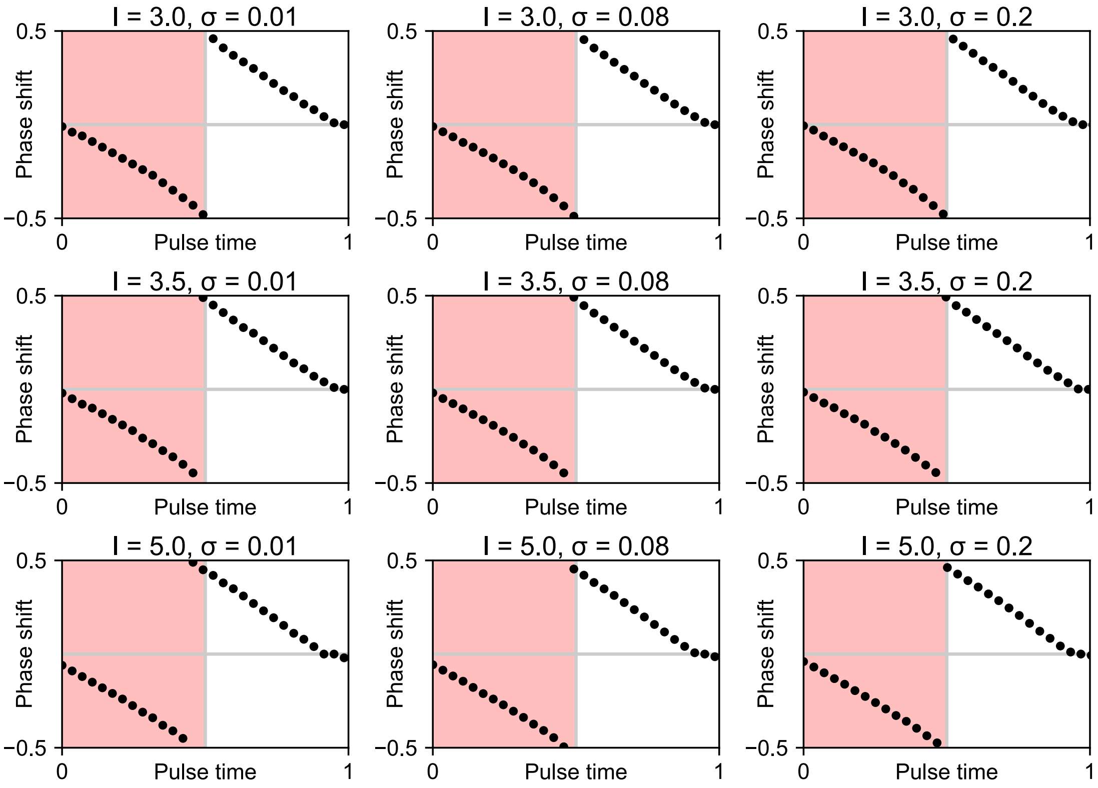


**Figure 16.** Phase response curves (PRCs) for varying noise intensity ($\sigma$) and input amplitude ($I$) for the noise-induced Van der Pol model. For the noise-induced model, the simulated PRCs have high amplitude regardless of noise intensity of input amplitude.
